# Photorespiration mediates plant carbon-nitrogen coordination

**DOI:** 10.64898/2026.11.22.768381

**Authors:** Naomi Nevo, Adi Yaaran, Moshe Alon, Efrat Sheffer, Yotam Zait

**Author notes:** Corresponding authors: Yotam Zait, <u>Email:</u>, Efrat Sheffer, <u>Email:</u>, Email addresses of all authors: Naomi Nevo,; Adi Yaaran,; Moshe Alon,; Efrat Sheffer,; Yotam Zait.

## Abstract

Plant species differ widely in their sensitivity to ammonium, and legumes are generally more sensitive than cereals, yet the mechanisms behind this difference remain unclear. Photorespiration links photosynthetic carbon assimilation with nitrogen metabolism, but its response to nitrogen form has rarely been compared between species. We grew the legume *Medicago truncatula*, with and without rhizobia, and the cereal wild emmer wheat (*Triticum dicoccoides*) under nitrate or ammonium, and measured gas exchange, chlorophyll fluorescence, leaf nitrogen, carbon isotope composition and amino acid profiles. Ammonium reduced biomass in both species, but through different leaf-level responses. In *M. truncatula*, ammonium increased leaf nitrogen but decreased CO_2_ assimilation, electron transport and photorespiratory CO_2_ release, lowering photosynthetic nitrogen-use efficiency. These effects were strongest in rhizobia-inoculated plants. In *T. dicoccoides*, ammonium increased photorespiration and photosystem II operating efficiency, photosynthesis and photosynthetic nitrogen-use efficiency were maintained, and nitrogen accumulated in asparagine and arginine-pathway amino acids. Photorespiration therefore responded in opposite directions to the same nitrogen form in the two species. We propose that sustained photorespiration helps maintain leaf carbon–nitrogen coordination under ammonium, whereas its suppression marks leaf-level ammonium sensitivity, with implications for efforts to engineer photorespiration in crops.

**Highlight:** Ammonium increased photorespiration and maintained photosynthetic nitrogen-use efficiency in wild emmer wheat but suppressed both in *Medicago truncatula*, most strongly in rhizobia-inoculated plants.

## Introduction

Improving photosynthesis has become a major target of crop breeding (Paul et al., 2026), motivated by the need to increase agricultural productivity for a growing global population and by recognition that further gains in yield are unlikely to stem solely from the morphological and biomass partitioning traits that formed the basis of the Green Revolution (Zhu et al., 2010). Future improvements, however, must occur in parallel with greater resource use efficiency, as continued increases in external inputs, particularly fertilizers, are not a sustainable long-term strategy (Hirel et al., 2007). Nitrogen is especially important in this context, as insufficient nitrogen availability can strongly limit plant growth and productivity, while supplied nitrogen is poorly utilized by crops (Sandhu et al., 2021). A large proportion of applied nitrogen is lost through leaching, runoff, and gaseous emissions, with significant environmental consequences (Li et al., 2017).

The need to reconcile high productivity with improved nitrogen use efficiency brings photosynthetic nitrogen use efficiency (PNUE) into focus. PNUE, typically defined as the rate of carbon assimilation per unit of leaf nitrogen, is particularly relevant because a large fraction of leaf nitrogen is invested in the photosynthetic apparatus (Evans and Clarke, 2019). Yet, high leaf nitrogen does not necessarily confer high photosynthetic capacity. The extent to which nitrogen investment is translated into carbon gain depends on its allocation and on the biochemical capacity for carbon assimilation, both of which vary across species and environments (Evans, 1989). Legumes, for example, often have high leaf nitrogen concentrations, without proportionally greater rates of carbon assimilation or growth compared with other C_3_ species (Wright et al., 2004; Adams et al., 2016; Bytnerowicz et al., 2023).

PNUE therefore reflects the coordination between carbon and nitrogen metabolism within the leaf, emphasizing the importance of the metabolic pathways that connect these processes (Lawlor, 2002).

Photorespiration is one such process, serving as a major metabolic link between photosynthetic carbon assimilation and nitrogen metabolism in illuminated C3 leaves (Eisenhut et al., 2019). It originates from the oxygenation reaction of Rubisco, producing 2-phosphoglycolate (2-PG), which is recycled through a series of reactions spanning the chloroplast, peroxisome, and mitochondrion (Bauwe, 2019). This cycle returns carbon to the Calvin-Benson cycle while generating glycine and serine, making photorespiration an important source of these amino acids in illuminated C3 leaves (Busch, 2020; Gashu et al., 2025). The mitochondrial conversion of glycine to serine releases CO_2_ and ammonia (NH_3_) which must be rapidly reassimilated via the glutamine synthetase–glutamate synthase GS/GOGAT pathway to prevent ammonium (NH_4_^+^) accumulation and maintain cellular nitrogen homeostasis (Bloom et al., 2010; Betti et al., 2014). Photorespiration also contributes carbon intermediates required for ammonium reassimilation and reducing equivalents that can support nitrate (NO_3_^-^) and nitrite (NO_2_^-^) reduction to ammonium (Busch et al., 2018).

Furthermore, PNUE may depend on nitrogen form and source, as nitrate and ammonium follow distinct assimilation pathways and place different demands on carbon metabolism and reducing power (Xu et al., 2012; Britto and Kronzucker, 2013; Bloom, 2015). Nitrate is relatively inert and can be stored in the vacuole (Granstedt and Huffaker, 1982). It is transported through the xylem to the shoot (Hachiya and Sakakibara, 2017), where it is sequentially reduced to nitrite in the cytosol and to ammonium in the chloroplast before incorporation in amino acids through the GS/GOGAT pathway (Fortunato et al., 2023). Nitrate reduction is strongly light dependent and requires substantial reducing power, with photorespiratory metabolism supplying part of this demand for reducing equivalents (Rachmilevitch et al., 2004; Nunes-Nesi et al., 2010).

In contrast, ammonium is highly reactive and its equilibrium with uncharged ammonia can disrupt cellular pH and ion homeostasis (Esteban et al., 2016). It must therefore be rapidly assimilated into organic compounds (Lancien et al., 2000), primarily in the roots through the GS/GOGAT cycle, with the resulting amino acids transported to the shoot (Hawkesford et al., 2012). Although ammonium assimilation bypasses the energy-intensive reduction steps required for nitrate assimilation (Britto and Kronzucker, 2002), it places a substantial demand on carbon metabolism by requiring carbon skeletons, together with reducing power, for amino acid synthesis. In roots, ammonium assimilation relies on NADH-dependent GOGAT, with reducing equivalents supplied by the oxidative pentose phosphate pathway (Guo et al., 2007). Despite its lower theoretical energetic cost, ammonium supplied as a sole nitrogen source often inhibits plant growth and photosynthesis, a response commonly referred to as ammonium toxicity (Liu and von Wirén, 2017; Wang, 2021). However, plant species have long been known to differ widely in their sensitivity to ammonium toxicity (Gerendás et al., 1997; Britto and Kronzucker, 2002; Esteban et al., 2016; Rivero-Marcos et al., 2024). Yet, after decades of research, there is still no clear explanation of the physiological mechanisms that determine this sensitivity and its variation among species (Kong et al., 2022; Xiao et al., 2023). In particular, it is unclear whether ammonium sensitivity originates in the leaf, where photosynthesis supplies the carbon and reducing power needed for nitrogen assimilation, or downstream of photosynthesis, in the allocation of carbon to nitrogen assimilation and growth.

Beyond inorganic nitrogen nutrition, symbiotic N_2_ fixation provides an additional route of nitrogen acquisition in some plants, primarily legumes. Within root nodules, rhizobia reduce atmospheric N_2_ to ammonium which is subsequently incorporated into organic nitrogen compounds and transported to the shoot (Tegeder and Masclaux-Daubresse, 2018). In return, the plant host supplies photosynthetically derived carbon, mostly in the form of sucrose, to sustain this energetically demanding process (Udvardi and Poole, 2013; Roy et al., 2020). This exchange represents another point of close integration between carbon and nitrogen metabolism.

Together, the contrasting demands associated with nitrate, ammonium and symbiotically fixed nitrogen on carbon metabolism and reducing power raise the question of how nitrogen form and source alter the integration of photorespiratory metabolism with these processes, and how this contributes to variation in PNUE and ammonium sensitivity. Such effects may involve changes in downstream photorespiratory metabolism, including carbon recycling, glycine-serine metabolism and ammonium reassimilation, or in photorespiratory flux itself. Although nitrate assimilation has been linked to photorespiration (Rachmilevitch et al., 2004; Bloom et al., 2010; Busch et al., 2018), photorespiratory responses to ammonium have rarely been compared between species that differ in ammonium sensitivity, or in legumes that also depend on symbiotic N_2_ fixation.

To address this question, we compared the legume *Medicago truncatula* (barrelclover) and the cereal *Triticum dicoccoides* (wild emmer wheat, accession Zavitan) under nitrate and ammonium nutrition. We chose these species because they belong to plant groups that differ in nitrogen acquisition strategy and in ammonium sensitivity. Legumes are generally more sensitive to ammonium than cereals (Rivero-Marcos et al., 2024), but the mechanisms behind this difference, and in particular how the uptake and assimilation of different nitrogen forms affect leaf carbon and nitrogen metabolism, have not been resolved. Comparing a species from a sensitive group with a species from a tolerant group allowed us to test how nitrogen form affects the coupling between nitrogen and carbon metabolism, and where along this pathway the two species diverge. Because external nitrogen supply can suppress symbiotic fixation (Zhong et al., 2023), we grew *M. truncatula* with and without external nitrogen, as well as with and without rhizobial symbionts to distinguish the effects of externally supplied nitrogen from those of nitrogen acquired by symbiotic N_2_ fixation.

We tested two alternative expectations, based on the contrasting demands of the two nitrogen forms for carbon and reducing power. If ammonium sensitivity arises mainly from the demand for carbon skeletons and energy for ammonium assimilation (Britto and Kronzucker, 2002; Vega-Mas et al., 2019), ammonium should reduce growth without impairing leaf photosynthesis. In this case, photorespiratory flux should be maintained or even increased under ammonium, because photorespiration supplies intermediates and reducing equivalents for nitrogen assimilation. Alternatively, if ammonium impairs leaf metabolism directly (Esteban et al., 2016; Liu and von Wirén, 2017), photosynthesis, electron transport and photorespiration should decline together, lowering PNUE despite high leaf nitrogen. We expected the first response in the more tolerant cereal and the second in the more sensitive legume. In *M. truncatula*, rhizobial symbiosis could either increase PNUE, because symbiotic sinks can stimulate photosynthesis (Kaschuk et al., 2010; Concha and Doerner, 2020), or decrease it, because nodules compete with ammonium assimilation for photosynthetic carbon delivered to the roots (Pate et al., 1979; Udvardi and Poole, 2013). Together, these comparisons allowed us to examine how photorespiratory metabolism contributes to variation in PNUE and ammonium sensitivity among nitrogen sources and species.

## Materials and Methods

### Plant Material

We grew wild emmer wheat (*Triticum turgidum* ssp. *dicoccoides* (Körn. ex Asch. & Graebn.) Thell., accession Zavitan) and barrelclover (*Medicago truncatula* Gaertn.) in 6 L pots filled with perlite (Perlite 2, Agrikal Ltd., Serik, Antalya, Turkey), an inert, pH-neutral medium with high water-holding capacity. We sowed wild emmer directly into pots. We surface-sterilized *M. truncatula* seeds in a chlorine solution for 2 min, rinsed them with ultrapure water, scarified the seed coat with sandpaper and germinated them on sterile agar before transplanting at the sprout stage.

One week after transplanting, we inoculated half of the *M. truncatula* plants with a mixture of *Sinorhizobium* strains previously isolated from nodules of naturally growing *M. polymorpha* collected at four geographically distinct sites in Israel. Strains were identified based on 16S rRNA sequencing and whole-genome analysis, and had been cultured on yeast extract mannitol (YEM) medium for 48 h (Alon et al., unpublished). We applied 1 mL of the bacterial suspension to the surface of the perlite at the base of each plant using a sterile pipette.

We conducted the experiment under net-house conditions (20% shading) at the Robert H. Smith Faculty of Agriculture, Rehovot, Israel, during the natural growing season (December-March). During the first two weeks, we irrigated all plants with a commercial fertilizer mixture containing NPK, Ca and Mg and micronutrients (Sheffer, Kalmagon and Kuratin; ICL Group Ltd., Tel Aviv-Yafo, Israel), supplying 100 ppm N to support early establishment. Thereafter, we irrigated the plants with full-strength Hoagland solution (see Supplementary Table S1) modified to provide nitrogen either as nitrate (KNO_3_) or ammonium ((NH_4_)_2_SO_4_) at a total concentration of 50 ppm N, allowing approximately 20% excess drainage. For *M. truncatula*, we included an additional nitrogen-free treatment in which nitrogen was omitted from the nutrient solution.

For *M. truncatula*, 15 replicate plants were included for each combination of nitrogen treatment and rhizobial inoculation (N=90). For *T. dicoccoides*, 15 replicate plants were included for each nitrogen treatment (N=30). To minimize rhizobial cross-contamination via irrigation, we positioned inoculated *M. truncatula* at the downstream end of the irrigation line, and placed *T. dicoccoides* between uninoculated and inoculated *M. truncatula* plants.

### Gas exchange and chlorophyll fluorescence measurements

We measured leaf gas exchange using the LI-6800 Portable Photosynthesis System (LI-COR Biosciences, Lincoln, NE, USA) equipped with a 2 cm² leaf chamber. We conducted measurements on the youngest fully expanded leaf between 08:00 and 14:00 at ambient temperature, with photosynthetically active radiation set at 1200 µmol photons m⁻² s⁻¹, a fan speed of 5000 rpm, and relative humidity maintained at 60%.

We obtained A/Ci response curves by sequentially adjusting reference CO₂ concentrations (µmol mol⁻¹) in the following order: 420, 300, 200, 100, 50, 420, 600, 800, 1000, 1200, 1500, and returning to 420. We performed light response measurements under both ambient (21% O₂) and low oxygen (1% O_2_ in air) conditions at a constant CO₂ concentration of 420 µmol mol⁻¹, using stepwise irradiances of 1200, 2000, 1800, 1600, 1400, 1200, 1000, 800, 600, 400, 300, 200, 100, and 50 µmol photons m⁻² s⁻¹.

We measured chlorophyll fluorescence simultaneously with the light response curves using the integrated pulse-amplitude modulation (PAM) fluorometer, which applied saturating pulses at each light step to determine steady-state (F_s_) and maximum fluorescence in the light adapted state (F_m_′).

### CO_2_ and light response curve fitting

We fitted A/Ci response curves to the Farquhar, von Caemmerer and Berry model of C₃ photosynthesis (Farquhar et al., 1980) to estimate the maximum rate of carboxylation (*V_cmax_*), the maximum rate of electron transport (*J_max_*) and day respiration (*R_d_*). We performed curve fitting using the fit_aci_response() function from the photosynthesis package (version 2.1.5) in R (Stinziano et al., 2021), using measured leaf temperature as model input. We retained the default assumption of infinite mesophyll conductance and thus interpreted V_cmax_ and J_max_ as apparent parameters that integrate biochemical limitation with the effects of mesophyll conductance.

We fitted light response curves using the Marshall and Biscoe model (Marshall and Biscoe, 1980), which describes the irradiance dependence of net photosynthesis in C₃ plants. To account for temperature variation during measurement, we standardized net assimilation rates to 25 ^°^C prior to fitting using an Arrhenius-type temperature response function (Bernacchi et al., 2001).

### Chlorophyll Fluorescence–Based Estimation of Photosynthetic Parameters

We used chlorophyll fluorescence measurements to assess photosystem II (PSII) photochemistry and to estimate total electron transport rate *(J_t_*), photorespiratory CO_2_ release (*F*) and the partitioning of electron flow between carboxylation ( *J_C_*) and oxygenation ( *J_O_*).

We calculated the effective quantum yield of PSII according to Genty et al. (1989) as:

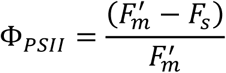

where *F*^′^ is the maximal fluorescence yield measured during a saturating pulse under light adapted conditions, and *F_s_* is steady state fluorescence yield in the light.

Fluorescence-based electron transport is commonly expressed as (Valentini et al., 1995):

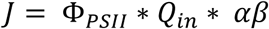

where *Q_in_* is incident photon flux density, α is leaf absorbance and β is the fraction of absorbed light partitioned to PSII. Rather than applying fixed values α and β, we calibrated Φ*_PSII_* against the quantum yield of CO_2_fixation (Φ*_CO_*_2_ ) following Valentini et al., 1995). At low O_2_, photorespiration is strongly suppressed and, assuming negligible alternative electron sinks, electron transport is primarily associated with CO_2_ fixation:

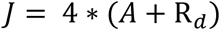

where A is the net CO_2_ assimilation rate, R*_d_* is day respiration, fixed at 1 µmol CO₂ m⁻² s⁻¹, and 4 is the number of electrons required per CO_2_ fixed.

Φ*_CO_*_2_ is defined as CO_2_ fixation per unit incident photon flux:

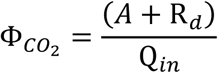

So that electron transport under low O_2_ can also be expressed as:

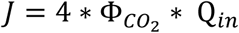

Using the 1% O_2_ data, we fitted the relationship between Φ*_PSII_* and Φ*_CO_*_2_ by linear regression (Supplementary Fig. S1):

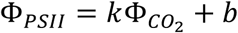

where *k* and *b* are the slope and intercept, respectively.

The fitted relationship was then used to calculate the apparent quantum yield of total linear electron transport:

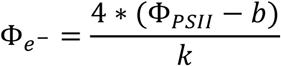

and total electron transport rate as:

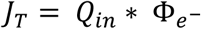

Following Cornic and Briantais (1991), we assumed *J_T_* to be predominantly partitioned between electron flux supporting RuBP carboxylation (*J_C_*) and oxygenation ( *J_O_*):

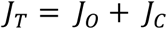

Electron flux associated with carboxylation was calculated as:

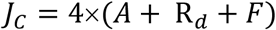

where F is the rate of photorespiratory CO_2_ release.

Assuming a photorespiratory stoichiometry of one CO_2_ released per two RuBP oxygenation reactions, *J_O_* can be calculated as:

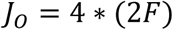

Combining these relationships gives:

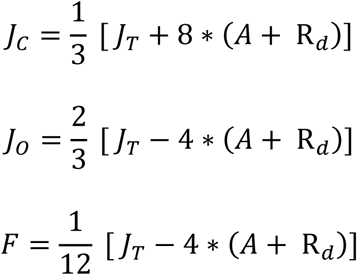

Net CO_2_assimilation used in this calculation was standardized to 25°C as described above. Regression-derived total electron transport was also standardized to account for temperature variation during measurements using an Arrhenius-type temperature response function (Bernacchi et al., 2003). The standardized values of *A* and *J_T_* were used to calculate *J_C_*, *J_O_* and *F*. Φ*_PSII_*, derived directly from chlorophyll fluorescence, was not temperature standardized.

### Biomass parameters

We harvested plants at the onset of inflorescence and separated roots and shoots, for all plants, and nodules for inoculated *M. truncatula* plants (uninoculated plants did not have any nodules). We oven-dried tissues at 60 °C, and then weighed them to determine dry mass.

### Chlorophyll quantification

At harvest we collected samples of fresh leaves and immediately froze them at −80°C until analysis. For pigment extraction, we added 500 µL of solvent to frozen leaf tissue and incubated the samples for 24 hours in the dark at room temperature. We used 100% ethanol as a solvent for *M. truncatula* and 80% acetone as a solvent for *T.* dicoccoides, as ethanol did not provide effective pigment extraction for this species. We measured absorbance at 665, 652, 470, and 750 nm using a microplate reader. We calculated chlorophyll and carotenoid concentrations according to Lichtenthaler and Buschmann (2001), using the equations specific to each solvent, and multiplied all values by a dilution factor calculated as the extraction volume divided by the dry mass of the extracted leaf tissue.

### Organic carbon and nitrogen concentrations

We ground oven-dried leaf samples to a fine powder. We measured organic carbon (C) and total nitrogen (N) from dried, ground leaf tissue using an elemental analyzer (Flash2000 CN Elemental Analyzer; Thermo Scientific, Milan, Italy).

### Stable carbon isotope composition (δ¹³C, ‰)

To determine stable carbon isotope composition, we weighed approximately 2–3 mg of dried ground leaf tissue and analyzed stable carbon isotope composition (δ¹³C, ‰) using a cavity ring-down spectroscopy (CRDS) system (G2131-I, Picarro Inc., Sunnyvale, CA, USA) calibrated with a cellulose standard, and expressed values relative to the Vienna Pee Dee Belemnite (VPDB) standard.

### Photosynthetic nitrogen use efficiency

We calculated PNUE as net CO₂ assimilation per unit leaf N on an area basis (Poorter and Evans, 1998). PNUE indicates how effectively leaf nitrogen is converted into instantaneous carbon gain (Poorter and Evans, 1998; Evans and Clarke, 2019). Because it is based on total leaf nitrogen, it also includes nitrogen not directly involved in photosynthesis, such as nitrate and free amino acids stored in the leaf. We first calculated treatment-mean leaf N per area (N_area_, g N m^−2^) from treatment-mean leaf mass per area (LMA, g m^−2^) and leaf N concentration measured by elemental analysis (% dry mass):

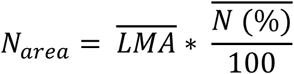

We then calculated PNUE for each replicate as:

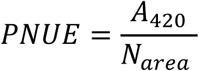

where *A*_420_ is the modeled net assimilation rate of individual samples at an intercellular CO₂ concentration of 420 µmol mol^−1^. PNUE is expressed as *μ*mol CO_2_ g^−1^N s^−1^ (Poorter and Evans, 1998). Because N_area_ was calculated from treatment means, within-treatment variation in PNUE reflects variation in A_420_ only.

### Symbiotic nitrogen fixation measures

At harvest time, we collected root nodules to estimate the momentary rate of biological nitrogen fixation by bacterial nitrogenase within the nodules, using the acetylene reduction assay (Hardy et al., 1968). We incubated a random sample of nodules from each plant in airtight jars under cool, dark conditions, and blank samples with no nodules. We replaced 10% of the air in each jar with acetylene following Dovrat et al. (2020). We quantified ethylene production after 30 and 60 minutes using a gas chromatograph equipped with a flame ionization detector (Focus; Thermo-Finnigan, Milan, Italy). The rate of activity within the nodules was calculated as amount of ethylene produced per gram of nodule mass per hour. We weighed all oven dried nodules for each plant and evaluated plant investment in symbiotic nitrogen fixation as the portion of biomass allocated to root nodules, calculated as total nodule mass per plant divided by total plant biomass.

### Analysis of amino acids

To evaluate amino acid profiles, we analyzed samples using an LC-MS system comprising a Nexera X2 system (Shimadzu, Kyoto, Japan) coupled to a QTRAP 6500+ mass spectrometer (SCIEX, Framingham, MA, USA). Chromatographic separation was achieved on a HILIC-Z column (150 × 2.1 mm, 3 μm; Agilent, Santa Clara, CA, USA) using a linear gradient of acetonitrile and water containing 100 mM ammonium formate. We operated the mass spectrometer in positive electrospray ionization mode and acquired data in multiple reaction monitoring mode.

### Statistical analysis

We performed statistical analyses in Python using the *pingouin* package (version 0.6.1). We treated individual plants as biological replicates. We assessed differences among treatments using Welch’s ANOVA which does not assume equal variances, followed by Games–Howell post hoc tests for pairwise comparisons. For A/Ci-derived parameters (*Vc_max_ , J_max_ , Vc_max_ / J_max_*), we analyzed each species separately, combining nitrogen source and inoculation status into a single treatment factor in *M. truncatula*, and using nitrogen source as the only factor in *T. dicoccoides.* For all other variables, we analyzed all treatment groups together, with each combination of plant species, nitrogen source, and inoculation status treated as a single treatment factor. We generated compact letter displays from the Games–Howell pairwise comparisons, with groups sharing a letter indicating that they did not differ significantly at P < 0.05. All tests were two-sided.

We identified outliers within each species × nitrogen × inoculation group using the interquartile range (IQR) method (1.5 × IQR), applied when group size was ≥ 4 (≥ 8 for δ¹³C). We applied this method to A/Ci derived parameters, δ¹³C and to ΦPSII-ΦCO₂ relationships measured under low-oxygen conditions.

## Results

### Biomass declines under ammonium in both species, but leaf nitrogen increases only in M. truncatula

Nitrogen form significantly affected total biomass, leaf nitrogen concentration and C:N ratio (P<0.001; Fig. 1a–c; Supplementary Table S2), with distinct responses across traits and species. In *M. truncatula*, ammonium-fed plants accumulated the highest leaf N concentration and had the lowest C:N ratios (Fig. 1b, c). In *T. dicoccoides* neither leaf nitrogen concentration nor C:N ratio differed significantly between nitrate and ammonium nutrition. However, under ammonium nutrition, *T. dicoccoides* tended to have lower leaf nitrogen than *M. truncatula*, whereas under nitrate nutrition it tended to have higher leaf N. These differences were significant only relative to uninoculated *M. truncatula*. Leaf chlorophyll concentration differed little among treatments despite differences in leaf N concentration, and was lowest in nitrogen-free, uninoculated *M. truncatula* (Fig. 1d). Biomass responded differently to nitrogen form (Fig. 1a). In both species, biomass was highest under nitrate nutrition and reduced under ammonium nutrition. Nitrogen-free, uninoculated *M. truncatula* plants had the lowest biomass, while rhizobial inoculation increased biomass under nitrogen-free conditions to levels comparable to ammonium-fed plants.

**Figure 1.**
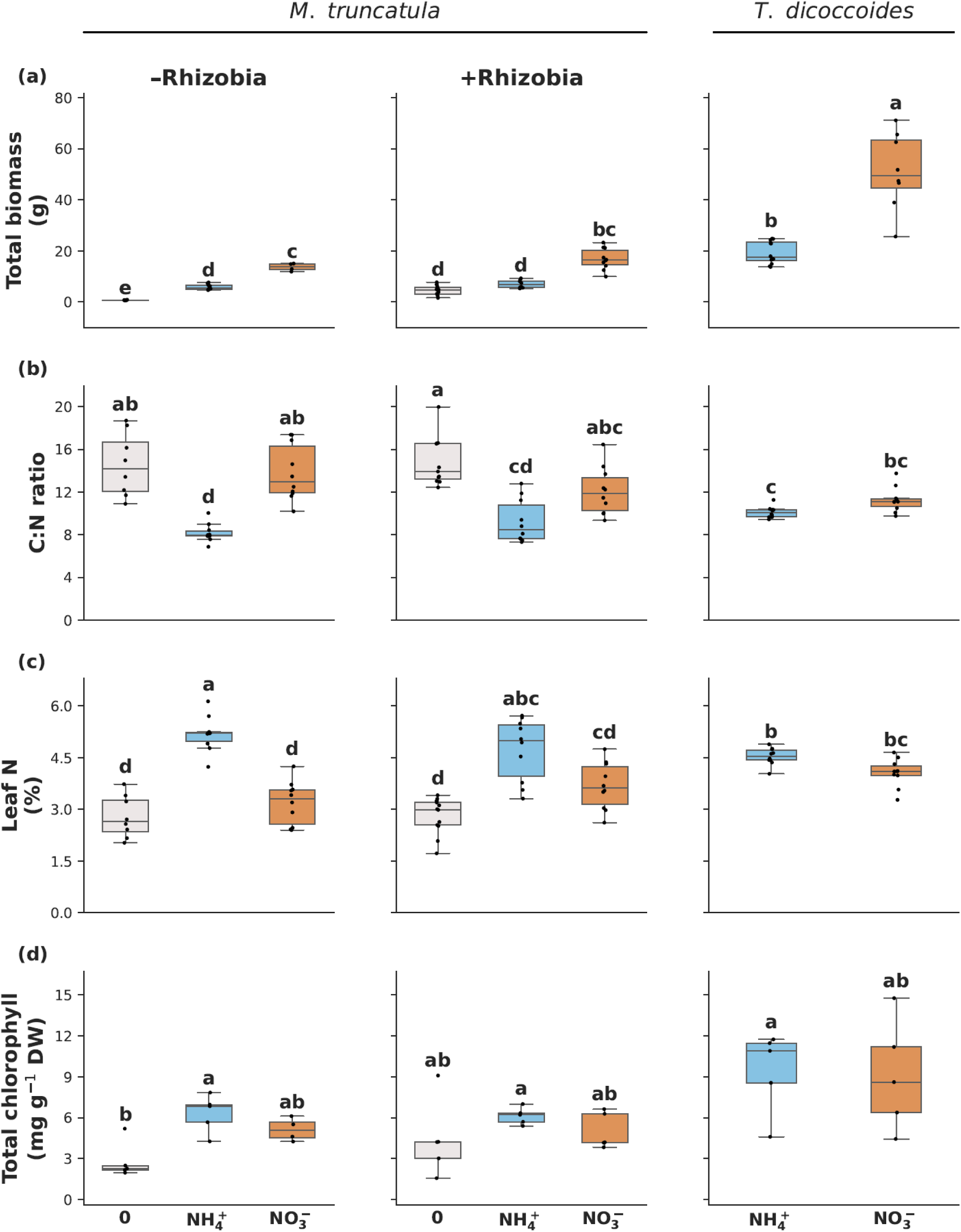
Effects of nitrogen form and rhizobial inoculation on growth and leaf traits. Dry weight biomass (a), leaf carbon-to-nitrogen (C:N) ratio (b), leaf nitrogen concentration (Leaf N%) (c), and total chlorophyll concentration (d) across treatments. Panels represent combinations of species (*Medicago truncatula* and *Triticum dicoccoides*), nitrogen source (nitrate, ammonium, or nitrogen-free), and inoculation status in *M. truncatula* (+Rhizobia, -Rhizobia). Boxplots show medians and interquartile ranges, with individual data points representing biological replicates (n= 5-8 per treatment). Different letters indicate significant differences among treatments (Welch’s ANOVA followed by Games–Howell post hoc test; P < 0.05). **Alt text:** Boxplots compare total dry biomass, leaf carbon-to-nitrogen ratio, leaf nitrogen concentration, and chlorophyll concentration among nitrogen treatments in *Medicago truncatula* and *Triticum dicoccoides.* Medicago treatments include plants with and without rhizobial inoculation. Points show individual plants, boxes show medians and interquartile ranges, and letters identify statistical groupings.

### Nitrogen supply alters nodule allocation and nitrogenase activity in M. truncatula

We next examined how external N supply affected plant investment in nodules and nitrogenase activity in inoculated *M. truncatula* (Fig. 2). The momentary rate of nitrogenase activity within the nodules was significantly higher in nitrogen-free plants, with no difference between the two external nitrogen treatments. Biomass allocation to nodules was highest under nitrogen-free conditions, intermediate under nitrate nutrition and lowest under ammonium nutrition.

**Figure 2.**
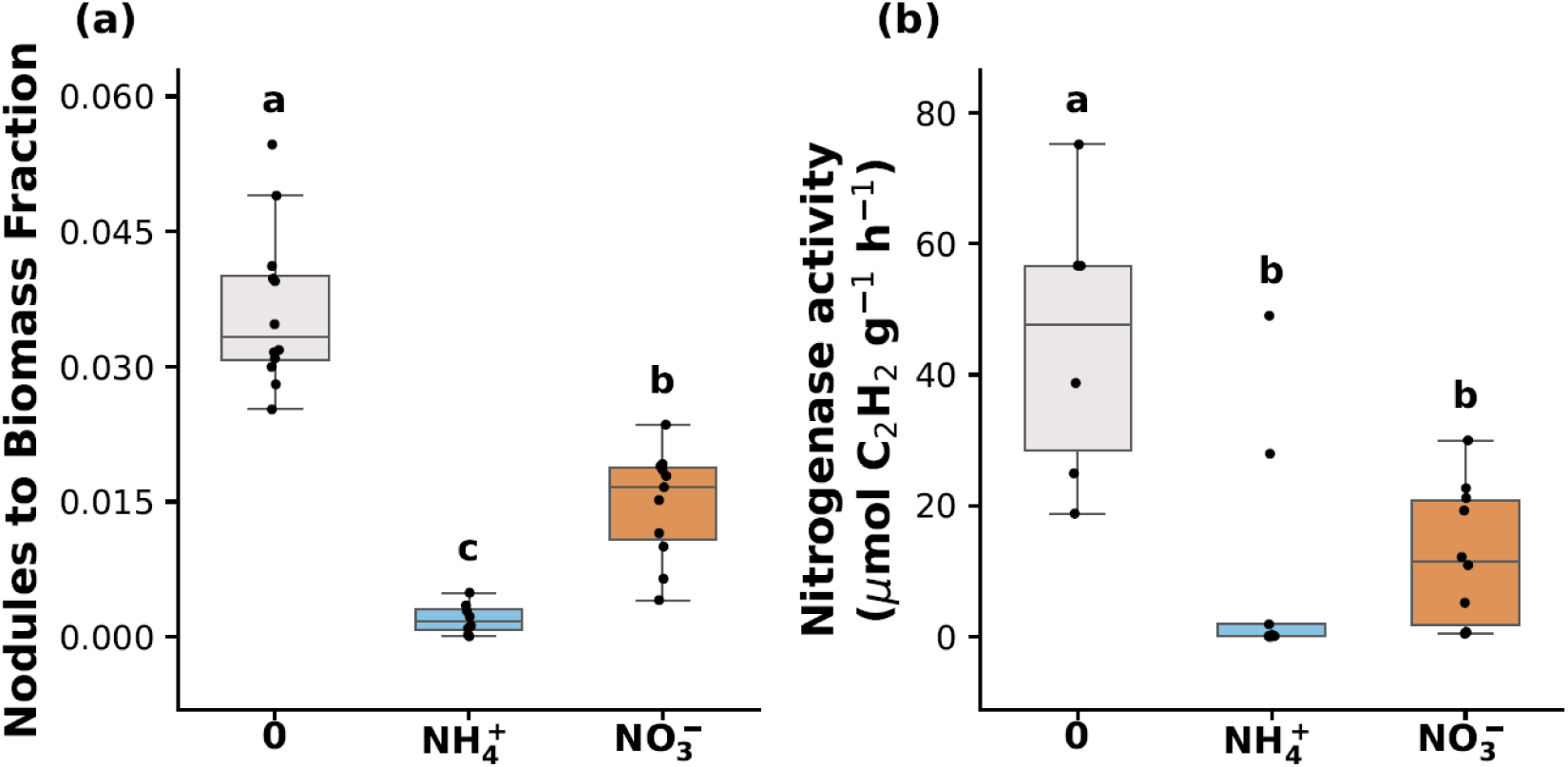
Symbiotic nitrogen fixation in *Medicago truncatula* under contrasting nitrogen sources. (a) Nodule biomass expressed as a proportion of total plant biomass in inoculated *M. truncatula* plants. (b) Nitrogenase activity within nodules, expressed as ethylene production rate per gram of nodule mass in the acetylene reduction assay. Boxplots show medians and interquartile ranges, with individual data points representing biological replicates (n ≥ 5 per treatment). Different letters indicate significant differences among treatments (Welch’s ANOVA followed by Games–Howell post hoc test; P < 0.05). **Alt text:** Two boxplots compare the fraction of plant biomass allocated to nodules and nodule nitrogenase activity in rhizobia-inoculated *Medicago truncatula* under nitrogen-free, nitrate, and ammonium nutrition. Individual points represent plants, and letters indicate statistical groupings.

### Ammonium inhibits photosynthesis in M. truncatula but not in T. dicoccoides

We assessed photosynthetic responses to nitrogen form using A/C_i_ curves (Fig. 3). In *M. truncatula*, CO_2_ assimilation was highest under nitrate nutrition and reduced under ammonium nutrition. Nitrogen-free, uninoculated plants had the lowest initial slope and maximum assimilation rate (*A_max_*). With rhizobial inoculation, the initial slope was similar to that of nitrate-fed plants, but Amax remained lower and comparable to inoculated ammonium-fed plants. Assimilation responses to nitrogen form differed with inoculation status. In uninoculated plants, differences between nitrate- and ammonium-fed plants decreased at high *C_i_*, but in inoculated plants they persisted across the *C_i_* range, higher in nitrate-than ammonium-fed plants. In *T. dicoccoides*, nitrate- and ammonium-fed plants differed only at low *C_i_*.

**Figure 3.**
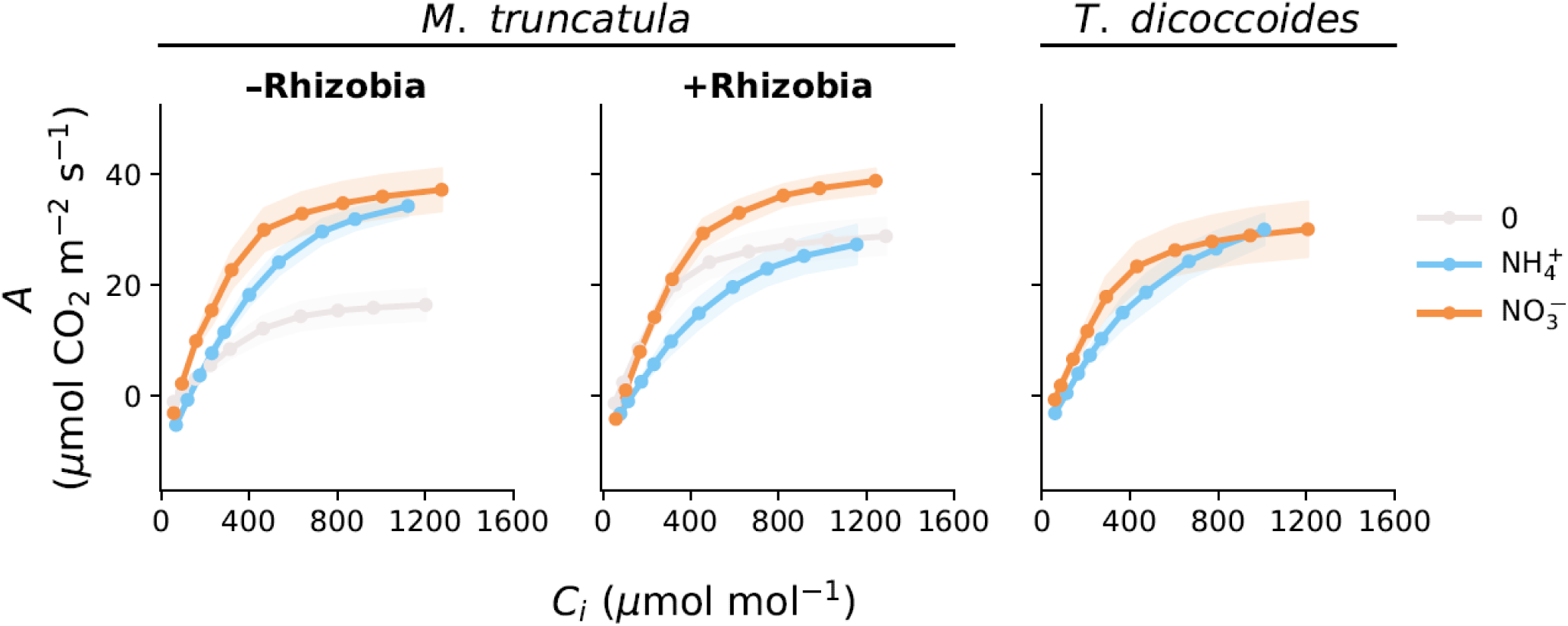
A/Ci response curves under different nitrogen sources and inoculation treatments. Net CO₂ assimilation rate (A) as a function of intercellular CO₂ concentration (Ci) for *M. truncatula* and *T. dicoccoides* grown under nitrate, ammonium, and (for *M. truncatula*) nitrogen-free conditions with or without rhizobial inoculation. Lines and symbols represent treatment means, and shaded areas denote ± SE (n ≥ 5). Curves were fitted using the Farquhar, von Caemmerer and Berry model of C_3_ photosynthesis. **Alt text:** Response curves show net carbon dioxide assimilation versus intercellular carbon dioxide concentration in *Medicago truncatula* and *Triticum dicoccoides* under different nitrogen treatments. Medicago plants are compared with and without rhizobial inoculation. Lines and symbols show treatment means, with shaded standard-error bands.

To further characterize photosynthetic biochemistry, we estimated the apparent maximum rate of Rubisco carboxylation (*V_cmax_*) and the maximum rate of electron transport (*J_max_*) from A/C_i_ curves (Fig. 4a–c; Supplementary Table S3). In *M. truncatula*, *V_cmax_* was lowest in nitrogen-free, uninoculated plants, significantly lower than in nitrate- and ammonium-fed uninoculated plants and nitrate-fed inoculated plants, and did not differ among the other treatments. *J_max_* varied with inoculation status (Fig. 4b). In uninoculated plants, *J_max_* was lower under nitrogen-free conditions than under ammonium nutrition, with nitrate-fed plants intermediate. In inoculated plants, *J_max_* was lower under ammonium than under nitrate nutrition, with nitrogen-free plants intermediate. The *V_cmax_/J_max_* ratio also depended on inoculation status. In uninoculated plants, it was highest under nitrogen-free conditions. In inoculated plants, the ratio was lower under nitrate than under ammonium nutrition, with nitrogen-free plants intermediate. In *T. dicoccoides*, neither *V_cmax_* nor *J_max_* differed between nitrate and ammonium nutrition, but the *V_cmax_/J_max_* ratio was lower under ammonium nutrition.

**Figure 4.**
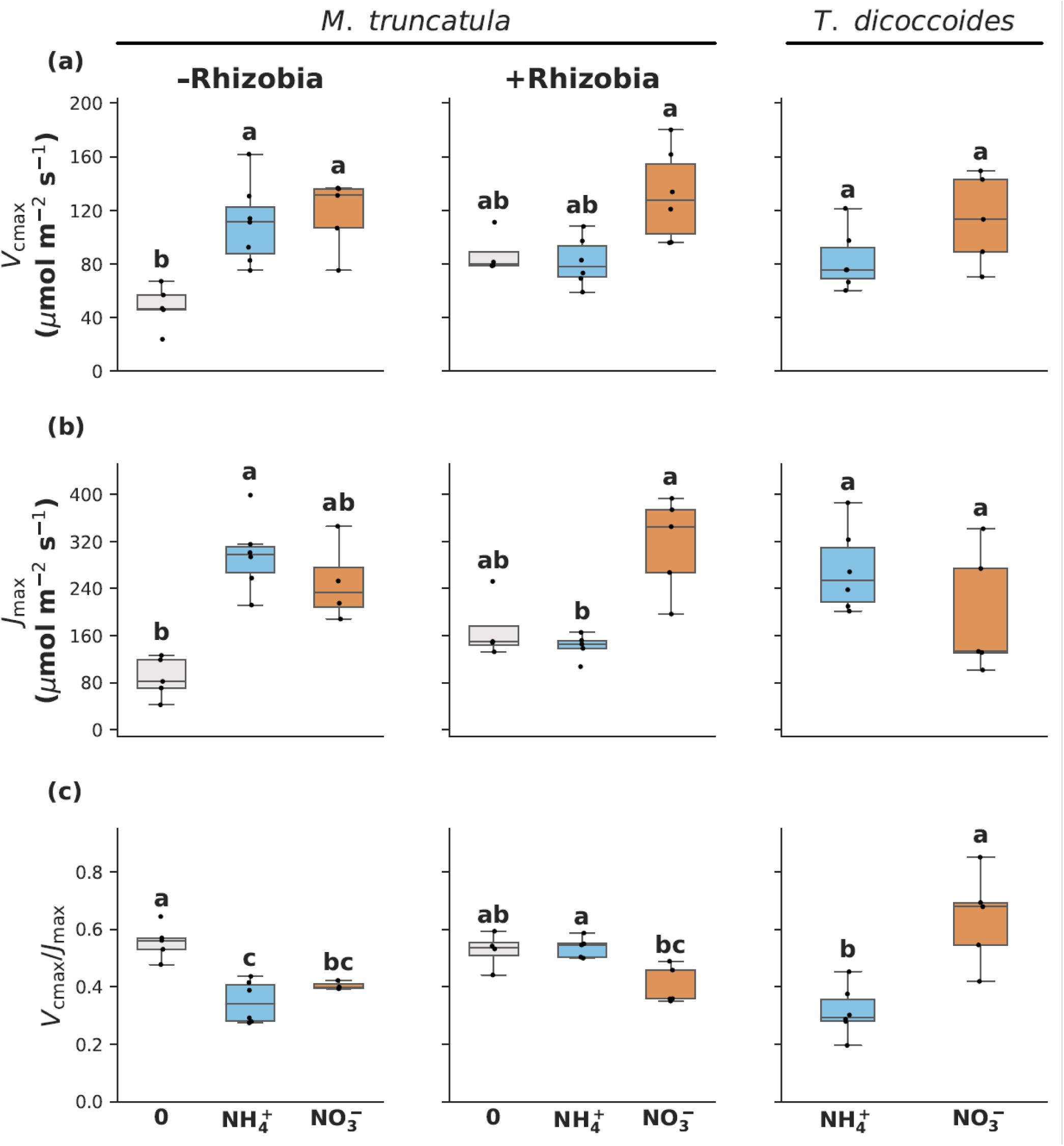
Photosynthetic parameters derived from A/Ci curves. (a) Maximum rate of carboxylation (V_cmax_), (b) maximum rate of electron transport (Jmax), and (c) V_cmax_/J_max_ ratio for *M. truncatula* and *T. dicoccoides* across nitrogen and inoculation treatments. +Rhizobia and −Rhizobia indicate inoculated and uninoculated *M. truncatula* plants, respectively. Boxplots show medians and interquartile ranges, with individual data points representing biological replicates (n ≥ 5 per treatment). Different letters indicate significant differences among treatments (Welch’s ANOVA followed by Games–Howell post hoc test; P < 0.05). **Alt text**: Boxplots compare apparent maximum Rubisco carboxylation capacity, maximum electron transport capacity, and their ratio across nitrogen treatments in *Medicago truncatula* and *Triticum dicoccoides*. Medicago groups also differ in rhizobial inoculation. Points show biological replicates, and letters indicate statistical groupings.

### Photorespiratory CO_2_ release increased under ammonium in T. dicoccoides but declined in M. truncatula

Photosynthetic responses to light were assessed using light-response curves (Fig. 5a), from which we derived electron transport rate (ETR; Fig. 5b) and photorespiratory CO_2_ release (F; Fig. 5c). In *M. truncatula*, assimilation across light levels was lower under ammonium than nitrate nutrition, with the largest difference in inoculated plants. In uninoculated plants, the two treatments had similar initial responses to irradiance, but assimilation remained lower under ammonium than under nitrate nutrition at high irradiance. Nitrogen-free inoculated plants had an initial response similar to that of nitrate-fed plants but reached lower assimilation at high irradiance. In *T. dicoccoides*, assimilation was similar under nitrate and ammonium nutrition.

**Figure 5.**
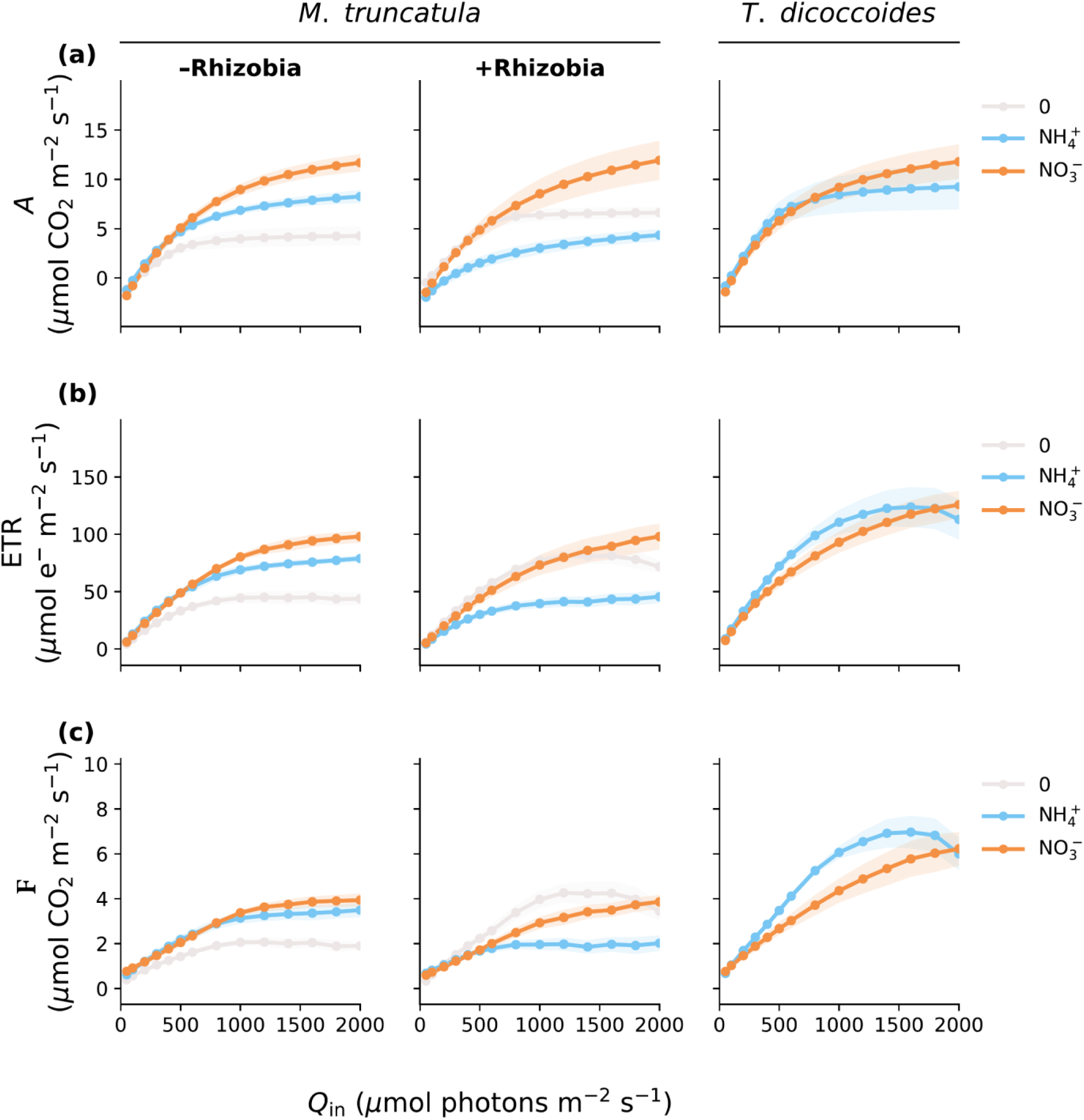
Light response of net assimilation, electron transport, and photorespiration. (a) Net CO₂ assimilation rate (A), (b) total electron transport rate (ETR), and (c) photorespiratory CO_2_ release rate (F) as functions of incident photon flux density (*Qin*) in *M. truncatula* and *T. dicoccoides* under the indicated nitrogen and inoculation treatments. Lines and symbols represent modeled means, and shaded areas indicate ± SE of the mean. **Alt text:** Light-response curves compare net carbon dioxide assimilation, total electron transport, and photorespiratory carbon dioxide release across nitrogen and inoculation treatments in *Medicago truncatula* and *Triticum dicoccoides*. Lines show treatment responses, with shaded standard-error bands.

ETR largely followed the assimilation response (Fig. 5b). At high irradiance, ETR was consistently higher in *T. dicoccoides* than in *M. truncatula* across all treatments, with the most pronounced difference under ammonium nutrition.

Photorespiratory CO_2_ release (*F*) was generally higher in *T. dicoccoides* than in *M. truncatula* (Fig. 5c), with contrasting responses to nitrogen form between the two species and between inoculation treatments in *M. truncatula*. In *T. dicoccoides*, *F* was higher under ammonium than nitrate nutrition across most of the light response curve, although this difference narrowed at high irradiance. In uninoculated *M. truncatula*, *F* was lowest under nitrogen-free conditions, while nitrate- and ammonium-fed plants had similar, higher values. In inoculated plants, nitrogen-free plants had the highest F over most of the irradiance range. Inoculated nitrate- and ammonium-fed plants were similar at low irradiance but diverged as irradiance increased, with *F* increasing under nitrate nutrition and remaining low under ammonium nutrition.

## PSII operating efficiency increases under ammonium in *T. dicoccoides* but declines in *M. truncatula*

The effective quantum yield of PSII (ΦPSII) responded differently to nitrogen form in the two species (Fig. 6). In *M. truncatula,* nitrate-fed plants had higher ΦPSII than ammonium-fed plants in both inoculation treatments. Among nitrogen-free plants, inoculated plants had the highest ΦPSII, while uninoculated plants approached nitrate-fed plants as irradiance increased. In contrast, in *T. dicoccoides* ΦPSII was higher under ammonium than nitrate nutrition, although this difference decreased at high irradiance.

**Figure 6.**
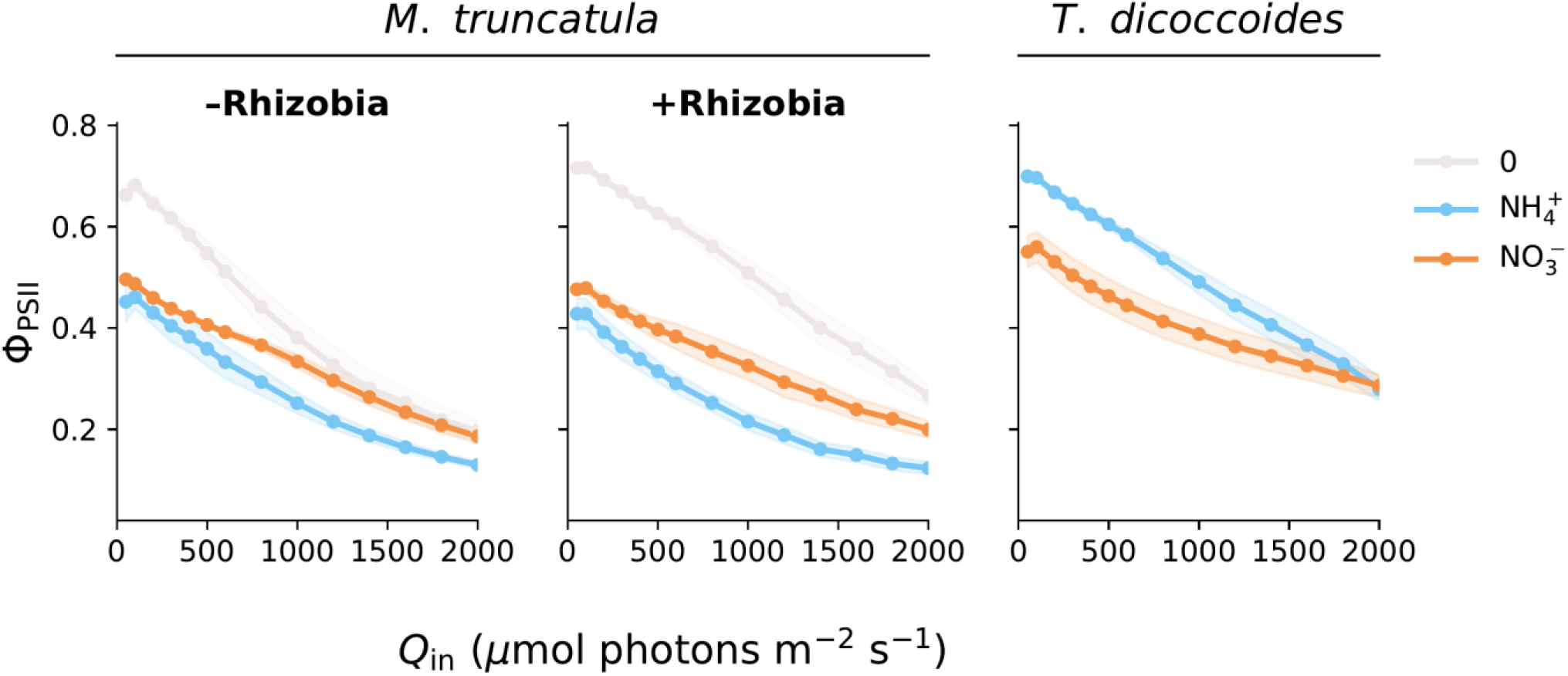
Light response of the effective quantum yield of photosystem II. Effective quantum yield of PSII (Φ_PSII_) as a function of incident photon flux density (*Qin*) in *M. truncatula* and *T. dicoccoides* under the indicated nitrogen and inoculation treatments. Lines and symbols represent treatment means, and shaded areas denote ± SE of the mean. **Alt text:** Light-response curves compare the effective quantum yield of photosystem II across nitrogen and inoculation treatments in *Medicago truncatula* and *Triticum dicoccoides.* Shaded bands represent standard errors.

### Carbon-isotope discrimination declines under ammonium, with a greater reduction in T. dicoccoides

Following the photosynthetic measurements, we examined leaf carbon isotope composition (δ¹³C; Fig. 7; Supplementary Table S4). In both species, δ¹³C was more negative under nitrate than ammonium nutrition. Under nitrate nutrition, values were similar between the two species, whereas under ammonium nutrition, *T. dicoccoides* had less negative δ¹³C than *M. truncatula*. In nitrogen-free plants, δ¹³C differed with inoculation status. Inoculated plants had the most negative values across treatments, while uninoculated plants were similar to nitrate-fed plants. Stomatal conductance was lower under ammonium than under nitrate nutrition in *M. truncatula* but changed little with nitrogen form in *T. dicoccoides* (Supplementary Fig. S2; Supplementary Table S5).

**Figure 7.**
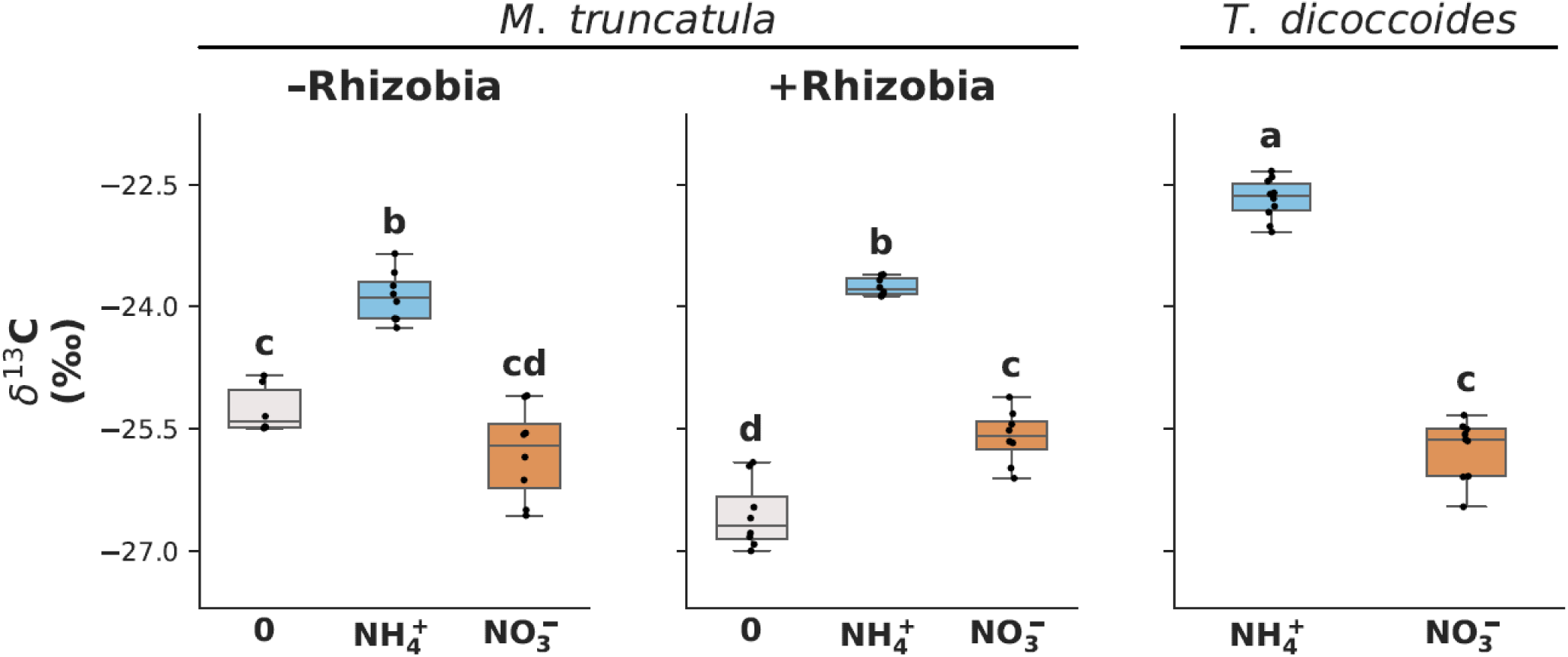
Leaf carbon isotope composition under contrasting nitrogen sources. Leaf carbon isotope composition (δ¹³C, ‰) of *M. truncatula* and *T. dicoccoides* grown under nitrate, ammonium, and (for *M. truncatula*) nitrogen-free conditions with or without rhizobial inoculation. Boxplots show medians and interquartile ranges, with individual data points representing biological replicates (n ≥ 5 per treatment). Different letters indicate significant differences among treatments (Welch’s ANOVA followed by Games–Howell post hoc test; P < 0.05). **Alt text:** Boxplots compare leaf carbon isotope composition across nitrogen treatments in *Medicago truncatula* and *Triticum dicoccoides,* including inoculated and uninoculated Medicago plants. Points show biological replicates, and letters identify statistical groupings.

### PNUE declines under ammonium despite higher leaf nitrogen in M. truncatula

We next compared photosynthetic nitrogen-use efficiency across nitrogen treatments and species (Fig. 8; Supplementary Table S6). In *M. truncatula*, PNUE was higher under nitrate than ammonium nutrition in both inoculated and uninoculated plants. Under nitrogen-free conditions, inoculation increased PNUE to values similar to those under nitrate nutrition. In *T. dicoccoides*, PNUE was similar under nitrate and ammonium nutrition and lower than in nitrate-fed *M. truncatula*.

**Figure 8.**
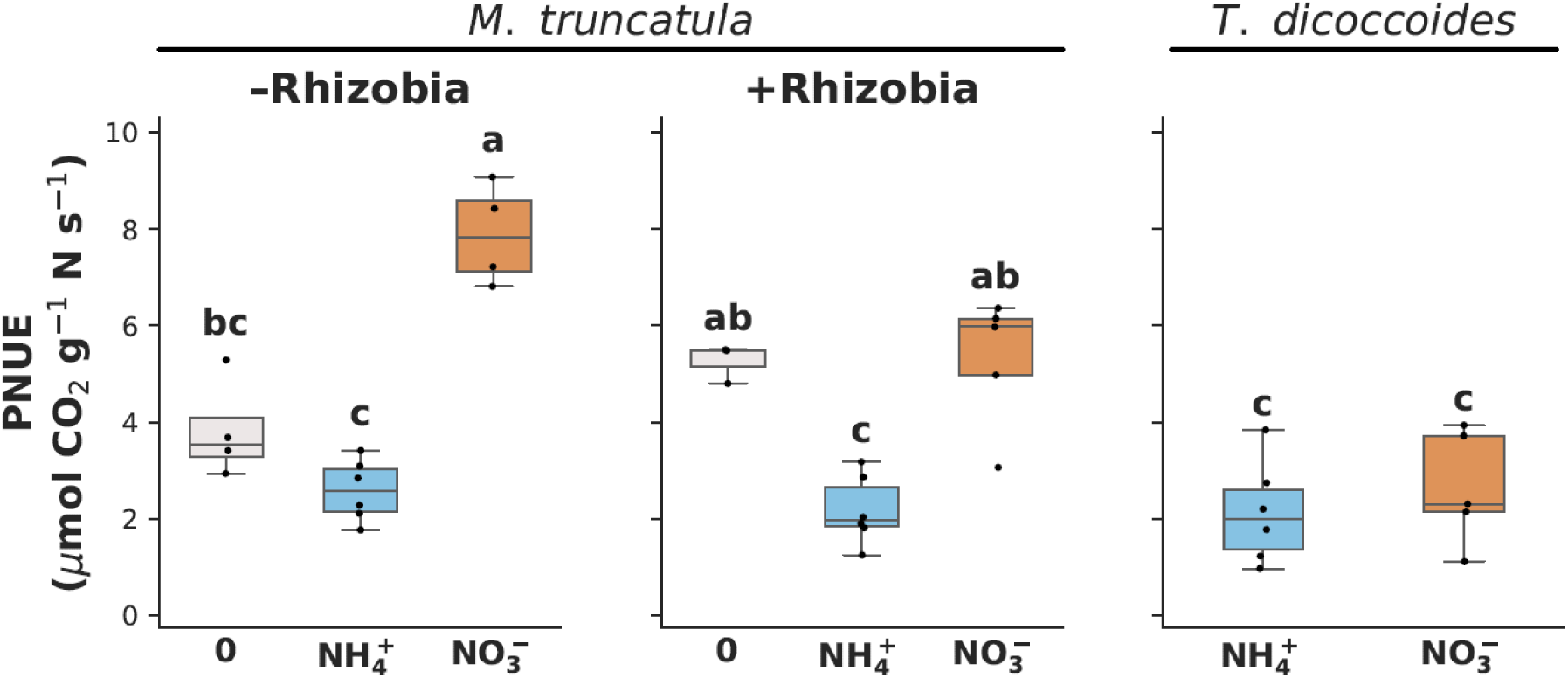
Photosynthetic nitrogen-use efficiency (PNUE). PNUE of *M. truncatula* and *T. dicoccoides* under the indicated nitrogen and inoculation treatments. PNUE was calculated as each replicate’s net assimilation rate predicted from its fitted A/Cᵢ curve at Cᵢ = 420 µmol mol⁻¹, divided by treatment-mean area-based leaf nitrogen content. Area-based leaf nitrogen was calculated from treatment-mean leaf mass per area and leaf nitrogen concentration (% dry mass). PNUE expresses carbon assimilation per unit nitrogen invested in the leaf and is reported in µmol CO₂ g⁻¹ N s⁻¹. Boxplots show medians and interquartile ranges, with individual data points representing biological replicates (n ≥ 5 per treatment). Different letters indicate significant differences among treatments (Welch’s ANOVA followed by Games–Howell post hoc comparisons; P < 0.05). **Alt text:** Boxplots compare photosynthetic nitrogen-use efficiency across nitrogen treatments in *Medicago truncatula* and *Triticum dicoccoides*, including Medicago plants with and without rhizobial inoculation. Points show biological replicates, and letters identify statistical groupings.

### Amino-acid profiles reveal different metabolic adjustments to ammonium in the two species

Finally, we examined leaf amino acid profiles across nitrogen treatments and species. Ammonium nutrition was associated with different amino acid profiles in the two species (Fig. 9; Supplementary Fig. S3). In *M. truncatula*, the relative abundance of glycine and serine was higher under ammonium nutrition than under nitrate nutrition or nitrogen-free conditions, in both inoculated and uninoculated plants. Additionally, the relative abundance of several other metabolites, including proline, hydroxyproline, histidine, citrulline and branched chain amino acids, was higher under ammonium nutrition, particularly in uninoculated plants. Glutamic acid followed a different pattern, with the highest relative abundance in uninoculated plants under nitrate nutrition. In *T. dicoccoides*, ammonium nutrition was associated with large increases in ornithine, agmatine, arginine and asparagine, and smaller increases in glutamine, glycine and serine, whereas proline and hydroxyproline did not change between nitrogen forms.

**Figure 9.**
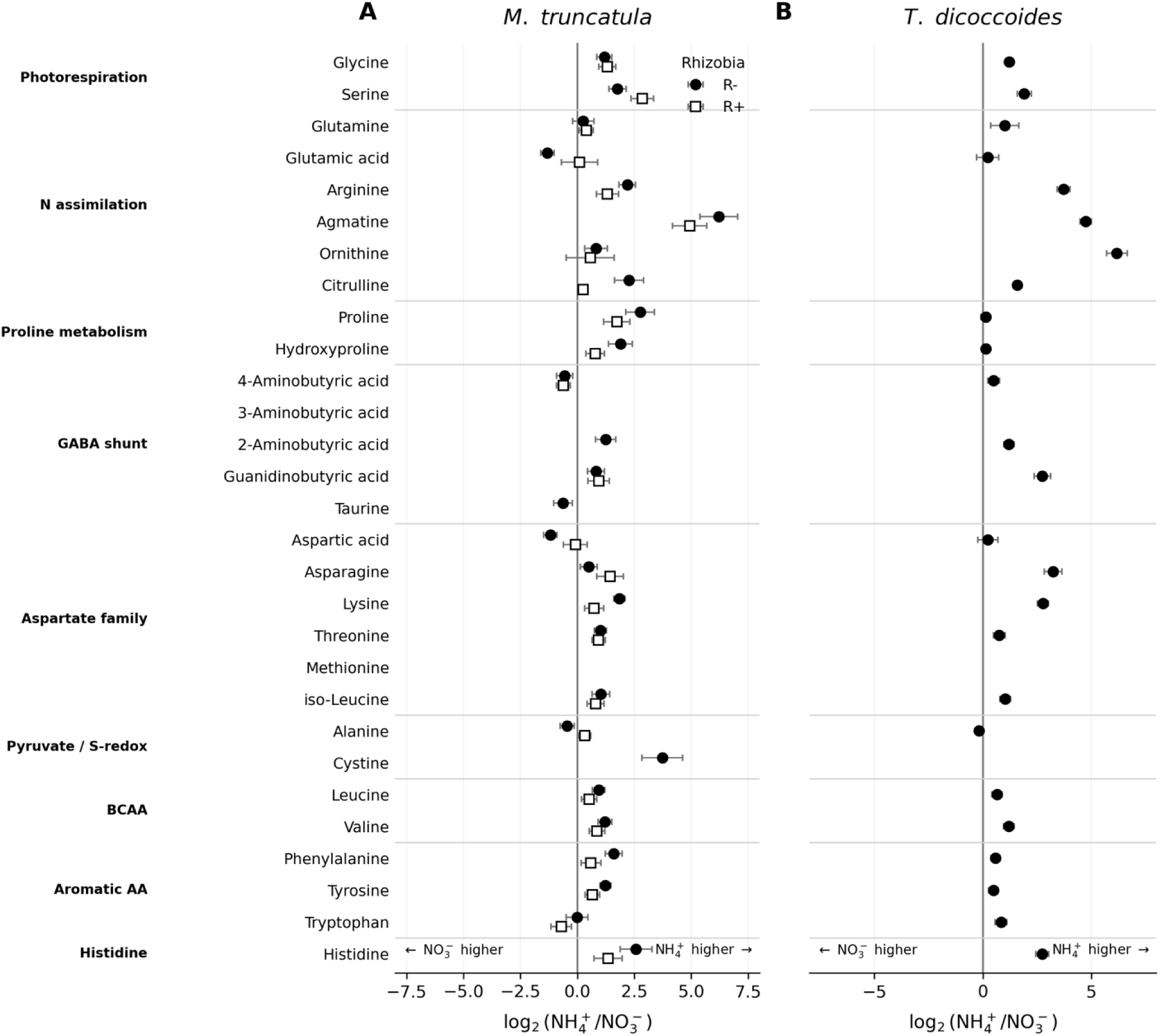
Leaf amino acid responses in *Medicago truncatula* and *Triticum dicoccoides*. Log₂ fold-changes in amino acid abundance under NH₄⁺ relative to NO₃⁻ are shown for *M. truncatula* (a) and *T. dicoccoides* (b). R+ and R− denote inoculated and uninoculated *M. truncatula*, respectively. Positive values indicate higher abundance under NH₄⁺, while negative values indicate higher abundance under NO₃⁻. Error bars represent propagated standard errors. Amino acids are arranged into functional groups: photorespiration-related amino acids; nitrogen assimilation; proline metabolism; γ-aminobutyric acid (GABA) shunt; aspartate metabolism; pyruvate-linked and sulfur/redox metabolism; branched-chain amino acids; aromatic amino acids; and the histidine family.

## Discussion

Our results show that nitrogen form affected photosynthetic nitrogen-use efficiency (PNUE) differently in *M. truncatula* and *T. dicoccoides*. In *M. truncatula*, ammonium nutrition substantially lowered PNUE relative to nitrate, while in *T. dicoccoides* PNUE was similar between the two nitrogen forms (Fig. 8). The basis of this contrast was apparent in the relationship between leaf nitrogen and carbon assimilation. Under ammonium nutrition, *M. truncatula* accumulated more leaf nitrogen but had lower photosynthetic rates, resulting in a significant decline in carbon gain per unit leaf nitrogen. In *T. dicoccoides*, by contrast, leaf nitrogen concentration and photosynthetic rates remained similar between ammonium- and nitrate-fed plants, and PNUE was consequently maintained (Figs 1c, 3, 5a). PNUE in *T. dicoccoides* was, however, lower than in nitrate-fed *M. truncatula* under both nitrogen forms, so the contrast between the species lies in the stability of PNUE rather than in its absolute value. Together, these results indicate that ammonium disrupted the coordination between nitrogen accumulation and carbon assimilation in *M. truncatula* but was largely maintained in *T. dicoccoides*. Biomass was nevertheless lower under ammonium than under nitrate nutrition in both species, despite these contrasting leaf-level responses, suggesting different processes underlying growth inhibition in each of the species (Fig. 1a). Ammonium sensitivity may therefore arise at different levels in the two species: in the photosynthetic machinery of the leaf in *M. truncatula*, and downstream of photosynthesis, in the use of carbon for nitrogen assimilation and growth, in *T. dicoccoides*.

The balance between carboxylation and electron transport (*V_cmax_/J_max_,* Fig. 4c) further distinguished the physiological response of the two species to ammonium. In *T. dicoccoides*, overall photosynthetic capacity changed little under ammonium, while the lower *V_cmax_/J_max_* ratio under ammonium reflected a shift to relatively greater electron transport capacity. In *M. truncatula,* the response depended on rhizobial inoculation. In uninoculated plants, *V_cmax_/J_max_* remained similar between nitrate and ammonium because *J_max_* (Fig. 4b) was maintained under ammonium, but in inoculated plants, the most natural condition for this species, ammonium reduced *J_max_* and increased the *V_cmax_/J_max_* ratio. The inoculation-dependent reduction in electron transport capacity was accompanied by a stronger inhibition of CO_2_ assimilation under ammonium (Figs 3, 4). Thus, in the legume, ammonium limited electron transport capacity, which supplies energy for both carbon fixation and photorespiration, whereas in the cereal, electron transport capacity was maintained relative to carboxylation capacity.

These adjustments in carboxylation and electron transport capacity were accompanied by distinct photorespiratory responses to ammonium nutrition in the two species. In *T. dicoccoides*, photorespiratory CO_2_ release (*F*; Fig. 5c) was higher under ammonium than nitrate nutrition, without a corresponding decline in CO_2_ assimilation or PNUE (Figs 5a, 8). In *M. truncatula*, by contrast, *F* declined under ammonium, particularly in inoculated plants, together with lower CO_2_ assimilation and PNUE. Electron transport rate (ETR; Fig. 5b) followed a similar overall pattern: in *T. dicoccoides* ammonium was associated with a modest increase in ETR at lower irradiance that diminished at high light, while in *M. truncatula* ETR was consistently lower under ammonium than nitrate nutrition, with the clearest reduction in inoculated plants. This correspondence between photorespiratory CO_2_ release and electron transport rate is consistent with the close energetic coupling between photorespiration and photosynthetic electron transport (Walker et al., 2024). Because photorespiration consumes ATP and reducing power, higher photorespiratory flux increases the demand for photosynthetically generated energy (Bauwe, 2023). This can provide an additional sink for electron transport and help prevent over-reduction of the photosynthetic electron transport chain, but it can also divert energy away from carbon fixation (Timm et al., 2024). Our results show that, in response to ammonium nutrition, higher photorespiratory flux in *T. dicoccoides* was associated with maintained CO_2_ assimilation and PNUE, whereas lower photorespiratory flux in *M. truncatula* coincided with photosynthetic decline. Under ammonium nutrition, photorespiration therefore did not behave as a simple carbon loss. Its maintenance accompanied a functional photosynthetic apparatus, consistent with the view that its net effect depends on how photorespiratory carbon and nitrogen are recycled or diverted into other pathways (Fu and Walker, 2023).

The chlorophyll fluorescence responses provide further insight into the photochemical context of the contrasting photorespiratory responses to ammonium (Fig. 6). In *T. dicoccoides*, ΦPSII remained higher under ammonium across most of the light response curve, indicating comparatively high PSII operating efficiency. In *M. truncatula*, ammonium consistently lowered Φ_PSII_, indicating reduced PSII photochemical performance. Together, these responses suggest that reduced photorespiration in *M. truncatula* formed part of a broader decline in PSII photochemical function, whereas in *T. dicoccoides* increased photorespiration coincided with higher PSII operating efficiency (Figs 5c, 6).

The species-dependent photochemical and photorespiratory responses to ammonium were also reflected in carbon isotope composition. In both species, δ¹³C was less negative under ammonium than nitrate nutrition, indicating lower carbon isotope discrimination, which was more pronounced in *T. dicoccoides* (Fig. 7). This stronger isotopic shift occurred despite little change in stomatal conductance (Supplementary Fig. S2), making stomatal limitation of CO_2_ diffusion an unlikely primary explanation and suggesting a greater contribution of non-stomatal processes to carbon isotope discrimination (Farquhar et al., 1989; Cernusak et al., 2013). Given the concurrent increase in photorespiratory CO_2_ release, changes in photorespiratory metabolism are one plausible contributor, although their specific effect on δ¹³C cannot be resolved from the present data. A second, non-exclusive explanation is that ammonium assimilation increases anaplerotic CO_2_ fixation by phosphoenolpyruvate carboxylase to supply carbon skeletons, which enriches plant tissue in ^13^C (Raven and Farquhar, 1990). This would be consistent with the larger investment in N-rich amino acids in *T. dicoccoides* (Fig. 9). In *M. truncatula*, by contrast, the less negative δ¹³C values occurred alongside lower stomatal conductance and reduced CO_2_ assimilation, suggesting a greater contribution of stomatal limitation and broader photosynthetic inhibition to the isotopic response (Lanigan et al., 2008).

The amino acid profiles provide a metabolic perspective on these physiological differences (Fig. 9). In *M. truncatula*, glycine and serine accumulated under ammonium despite lower photorespiratory CO_2_ release. Because their pool sizes reflect both production and downstream metabolism, this pattern is more consistent with altered turnover or use of photorespiratory metabolites than with increased Rubisco oxygenation (Fu et al., 2022). Proline and hydroxyproline also increased under ammonium, particularly in uninoculated plants, indicating a substantial stress-associated metabolic adjustment (Krasensky and Jonak, 2012; Verslues, 2024). Notably, this stronger metabolite accumulation occurred in uninoculated plants, which also had less photosynthetic inhibition. The weaker response in inoculated plants may therefore reflect tighter carbon and energy constraints associated with the additional cost of symbiosis (Pate et al., 1979; Taylor and Menge, 2018). *T. dicoccoides* had a different amino-acid profile. Glycine and serine increased to a similar extent as in *M. truncatula*, consistent with the higher photorespiratory flux, but proline and hydroxyproline did not change. Instead, ammonium nutrition strongly increased ornithine–arginine pathway metabolites and asparagine, with a smaller increase in glutamine, consistent with greater incorporation of nitrogen into N-rich amino acid pools (Vega-Mas et al., 2019). This greater investment in nitrogen assimilation may have increased the demand for carbon skeletons and reducing power, providing a possible explanation for the substantial biomass decline despite relatively stable photosynthesis and PNUE (Setién et al., 2013; Poucet et al., 2021).

Together, our results support the hypothesis that nitrogen form and source influence PNUE through their effect on photorespiration and its coordination with carbon and nitrogen metabolism (Fig. 10). In *T. dicoccoides*, ammonium was associated with increased photorespiratory CO_2_ release while carbon assimilation and PNUE were maintained, suggesting that photorespiration may buffer the altered carbon and energetic demands imposed by ammonium nutrition. The increase in glutamine and other N-rich metabolites further suggests that carbon and energy remained available to support nitrogen assimilation, although this was not sufficient to prevent substantial biomass reduction. In *M. truncatula*, by contrast, ammonium suppressed photorespiration together with carbon assimilation and PNUE. The accumulation of glycine and serine, alongside stress-associated metabolites such as proline and hydroxyproline, points to altered photorespiratory metabolite processing and a broad stress response. The suppression of photorespiration and carbon assimilation was strongest in inoculated plants, consistent with tighter carbon and energetic constraints associated with symbiosis.

**Figure 10.**
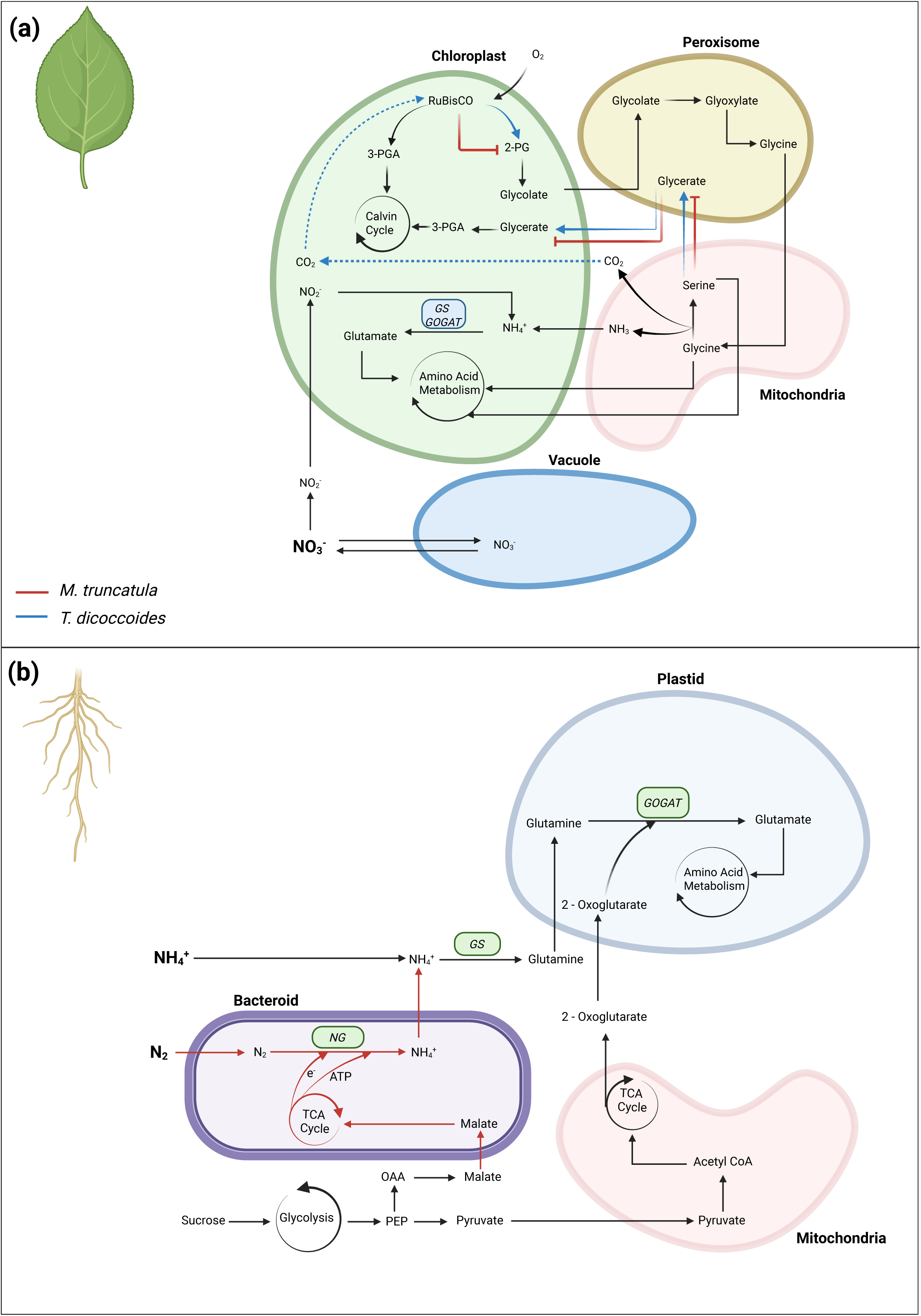
Conceptual model of nitrogen-form effects on photorespiration and PNUE in *Medicago truncatula* and *Triticum dicoccoides*. (a) In the leaf, xylem-derived nitrate (NO₃⁻) is either stored in the vacuole or reduced in the cytosol to nitrite (NO₂⁻), which is transported into the chloroplast, reduced to ammonium (NH₄⁺) and assimilated through GS-GOGAT. Photorespiration proceeds across the chloroplast, peroxisome and mitochondrion. Mitochondrial glycine-to-serine conversion releases CO₂ and NH₃; NH₃ is largely reassimilated through chloroplastic GS-GOGAT, whereas some glycine and serine enter broader amino-acid metabolism. The remaining photorespiratory carbon returns to the Calvin– Benson cycle as glycerate and 3-phosphoglycerate. Photorespired CO₂ may diffuse from the leaf or return to the chloroplast for refixation by Rubisco. Red inhibition symbols indicate processes proposed to be suppressed in *M. truncatula*, whereas blue arrows indicate processes proposed to be enhanced in *T. dicoccoides*. The dotted blue arrow denotes the proposed return of photorespired CO₂ to the chloroplast in *T. dicoccoides*. These responses are inferred from the present physiological and metabolic measurements and do not represent directly measured fluxes. (b) In the root, ammonium (NH₄⁺) taken up through transporters is assimilated through cytosolic GS and plastidial GOGAT, producing glutamate for further amino-acid metabolism. Phloem-delivered photoassimilates, primarily sucrose, support this process by supplying energy and carbon skeletons through glycolysis and mitochondrial TCA-cycle metabolism, including the 2-oxoglutarate required for GOGAT. In nodulated *M. truncatula*, sucrose-derived carbon is also supplied to bacteroids as malate, whose metabolism supports respiration and provides the reducing equivalents and ATP required for nitrogenase-dependent reduction of atmospheric N₂ to NH₃/NH₄⁺. Fixed nitrogen is subsequently released into the plant compartment and assimilated through the GS–GOGAT pathway. Red arrows indicate pathways associated with *M. truncatula*. **Alt text:** Conceptual diagram links nitrogen assimilation, photorespiration, and carbon metabolism in leaves and roots. Leaf pathways span chloroplasts, peroxisomes, mitochondria, and the cytosol; root pathways include ammonium assimilation and rhizobial nitrogen fixation. Red inhibition symbols mark proposed suppression in *Medicago truncatula*, and blue arrows mark proposed enhancement in *Triticum dicoccoides*. A dotted blue arrow indicates proposed refixation of photorespired carbon dioxide. These pathways represent inferred mechanisms rather than directly measured fluxes

Our results therefore suggest that photorespiration acts as a buffer for carbon–nitrogen coordination under ammonium, helping to accommodate the associated changes in energetic demand. This buffering was maintained in *T. dicoccoides* but disrupted in *M. truncatula*, where ammonium uncoupled nitrogen accumulation from carbon gain and led to broader metabolic impairment.

To our knowledge, this is the first study to compare photorespiratory responses to nitrate and ammonium between a legume and a cereal, with and without rhizobial symbiosis. It shows that photorespiration responds in opposite directions to the same nitrogen form in the two species, and that this divergence tracks PNUE. It also shows that ammonium inhibition of photosynthesis in the legume is strongest when plants rely on rhizobia, the natural state of legumes in the field, and that ammonium inhibits growth in both species through different physiological routes. These findings suggest that the long-recognized difference in ammonium sensitivity between legumes and cereals may arise, at least in part, from how photorespiration and photosynthetic electron transport respond to ammonium in the leaf. They also suggest that photorespiration should not be viewed only as a carbon loss to be minimized, as in current efforts to engineer photorespiratory bypasses (South et al., 2019), but also as a pathway whose capacity may need to be maintained when crops rely on ammonium-based fertilizers.

Our conclusions are based on one genotype of each species and on photorespiratory rates estimated from gas exchange and chlorophyll fluorescence rather than measured directly. Comparisons across more species pairs, together with isotopic flux analysis of photorespiratory metabolism (Fu et al., 2022; Gashu et al., 2025), are needed to test how general these responses are.

### Supplementary data

The following supplementary data are available at *JXB* online.

Table S1. Composition of the modified Hoagland solutions used for the nitrogen treatments.

Fig. S1. Relationship between PSII quantum yield and CO_2_ fixation quantum yield under low-oxygen conditions.

Table S2. Welch’s ANOVA results for biomass, leaf carbon-to-nitrogen ratio, leaf nitrogen concentration, and chlorophyll concentration.

Table S3. Welch’s ANOVA results for photosynthetic parameters derived from A/C_i_ curves. Table S4. Welch’s ANOVA results for leaf carbon isotope composition (δ^13^C).

Fig. S2. Stomatal conductance to water vapour across nitrogen and inoculation treatments. Table S5. Welch’s ANOVA results for stomatal conductance to water vapour.

Table S6. Welch’s ANOVA results for photosynthetic nitrogen-use efficiency.

### Fig. S3. Heat map of relative leaf amino acid abundance across nitrogen and inoculation treatments. Acknowledgements

We thank Julius Ben-Ari and the staff of the ZABAM Interdepartmental Equipment Unit in HUJI for assistance with LC-MS measurements. We thank Tania Masci for running all other laboratory analyses, Amit Sivan, Dvir Mendelson, Bar Ben Zeev and Ariel Joseph for their help in the experiment.

### Author contributions

NN: methodology, investigation, formal analysis, writing – original draft; AY: formal analysis, writing – review and editing; MA: methodology, resources; ES and YZ: conceptualization, methodology, supervision, writing – review and editing.

### Data availability

All code and data supporting the findings of this study are openly available on GitHub at https://github.com/zaitlab/Photorespiration-influences-photosynthetic-nitrogen-use-efficiency-across-different-plant-groups.

## Abbreviations

A: net CO_2_ assimilation rate
C_i_: intercellular CO_2_ concentration
ETR: electron transport rate
F: rate of photorespiratory CO_2_ release
GS/GOGAT: glutamine synthetase/glutamate synthase
J_C_: electron flux to RuBP carboxylation
J_max_: maximum rate of electron transport
J_O_: electron flux to RuBP oxygenation
J_T_: total electron transport rate
PNUE: photosynthetic nitrogen-use efficiency
PSII: photosystem II
R_d_: day respiration
RuBP: ribulose-1,5-bisphosphate
V_cmax_: maximum rate of Rubisco carboxylation
Φ_CO2_: quantum yield of CO_2_ fixation
Φ_PSII_: effective quantum yield of PSII.

